# Mast cell extracellular granules are bioactive condensates driven by heparin and polyamine

**DOI:** 10.64898/2025.12.25.696514

**Authors:** Yiwei Xiong, Dylan T. Tomares, Jianjian Guo, Kazuki Sato, Longhui Zeng, Yuan Tian, Maohan Su, Ava Albis, Avnika Pant, Rohit V. Pappu, Xiaolei Su

**Affiliations:** Department of Cell Biology, Yale School of Medicine, New Haven, CT 06520, USA; Department of Biomedical Engineering, Center for Biomolecular Condensates, James McKelvey School of Engineering, Washington University in St. Louis, St. Louis, MO 63130, USA; Yale Cancer Center, New Haven, CT 06520, USA; Yale Center for Immuno-Oncology, New Haven, CT 06520, USA; Yale Center for Systems and Engineering Immunology, New Haven, CT 06520, USA; Yale Stem Cell Center, New Haven, CT 06520, USA

## Abstract

Biomolecular condensates are membraneless bodies that organize biochemical reactions typically within cells. However, the roles of condensates in extracellular space—where conditions differ substantially from intracellular space—remain poorly understood. Here, we report mast cell extracellular granules (MCEGs), a stable membraneless entity, are condensates assembled via electrostatic interactions between glycosaminoglycans and polyamines. Disrupting polyamine synthesis or trafficking blocks MCEG formation and compromises the storage of proteases and cytokines. Granules reconstituted with heparin and spermine are sufficient to enrich mediators such as CPA3 and TNFα, maintaining an elevated pH and higher concentrations of calcium and zinc compared to the extracellular milieu. This unique environment enhances CPA3 enzymatic activity. Furthermore, the granules increase TNFα binding and its bioactivity toward endothelial cells. Together, we reveal MCEGs as functionally active biomolecular condensates with distinct biochemical and immunological properties; MCEGs are formed through sugar-metabolite interactions, expanding the mechanisms of condensate assembly beyond classical protein-protein and protein-RNA interactions.

## INTRODUCTION

Biomolecular condensation governs the spatial and temporal organization of the intracellular space by forming membraneless structures such as nucleoli^1–4^, nuclear speckles^5^, stress granules^6–8^, and immune signaling clusters^9–14^. Condensation can be mechanistically realized via a combination of reversible site-specific binding, percolation involving hierarchies of multivalent homotypic and heterotypic interactions, and solubility considerations that drive phase separation^1, 10, 15–18^. However, two major aspects of condensation have not received much attention. First, with a few exceptions^19–22^, the possibility of condensation playing a role in spatial and temporal organization of macromolecules and biochemical reactions in extracellular spaces has not been considered. Second, in cells and tissues, complex coacervation, as opposed to phase separation driven mainly by homotypic interactions among intrinsically disordered proteins, is likely to be the most important contributor to phase separation in multicomponent systems comprising highly-charged molecules that form compositionally distinct or multiphase biomolecular condensates^8, 23–38^. Complex coacervation is driven by system-specific and solution condition-dependent hierarchies of heterotypic interactions^25–27, 31–33, 39, 40^. The extracellular space has a distinct chemical environment in terms of its pH and compositions of ions and metabolites, which indicates a separate set of mechanisms for complex coacervation. Here, we present results from studies of extracellular granules of mast cells^41^ as archetypal systems to explore how condensation influences spatial organization and functions of biomolecules in the extracellular space.

Mast cells are long-lived immune cells that reside in tissues close to host-environment interfaces, such as the skin, intestinal, and pulmonary mucosa^42^. They are well known for their innate capabilities as responders to allergic inflammation and pathogen infections^43, 44^. There is also growing interest in the roles of mast cells in host-protective behavior through crosstalk with neurons^45–47^. Central to these versatile functions are mast cell granules, which are membrane-bound intracellular vesicles that serve as storage reservoirs of preformed bioactive mediators, including cytokines, chemokines, proteases, lipids, biogenic amines, and proteoglycans^48^. Upon activation by allergens, pathogens, or environmental stimuli, intracellular granules undergo exocytosis, during which the membrane coat is shed, and stored mediators are released into the extracellular space^49, 50^. While some mediators, such as histamine, are released in a soluble form, other mediators, including Tumor necrosis factor alpha (TNFα)^41^, interleukin-1 beta (IL-1β)/pro-IL-1β ^51^, tryptase^52^, chymase, and heparin proteoglycans^53, 54^ remain embedded in the membrane-free “granule remnants”^55^. These stable mast cell extracellular granules (MCEGs) are micron-sized bodies and are readily observed under the microscope^56, 57^. These intact MCEGs not only modulate macrophage and dendritic cell functions locally^58, 59^, but also enter the blood and lymphatic vessels to induce immune response remotely^41, 60^. However, the molecular mechanism for maintaining these membraneless MCEGs in the extracellular space remains unclear. The lack of this knowledge also impedes understanding of the benefits of maintaining mediators in the granule versus in the soluble format.

Heparin and chondroitin sulfate glycosaminoglycans are the major components of mast cell granules^48, 61^. Heparin is synthesized selectively in mast cells^48^. Genetic deletion of key heparin-synthesizing enzymes like N-deacetylase/N-sulfotransferase 2 (Ndst2) or glycosyltransferase Exostosin 1 impairs granule formation^62–64^, highlighting the essential role of heparin in granule assembly. Further, because of the highly negative charge of heparin^65^, positively charged cargo proteins may thus complex with heparin via electrostatic interactions. Recent studies found that cell surface heparan sulfate promotes the condensation of highly basic proteins such as basic fibroblast growth factor (bFGF)^66^ and CCL5^22^. However, these proteins are not the major components of MCEGs. Nevertheless, the studies on condensates formed by heparan sulfate and bFGF / CCL5 inspired us to test the possibility that the negatively charged heparin can undergo coacervation with positively charged biomolecules in MCEGs to drive condensate formation, thus providing the structural basis for the assembly of membraneless MCEGs.

Here, we report that MCEGs are formed through the condensation between heparin and polyamine. Using reconstitution approaches, we showed that condensates containing heparin and spermine create a distinct electrochemical microenvironment characterized by elevated pH and enriched metal ions. Notably, this specialized environment enhances the enzymatic activity of protease carboxypeptidase A3 (CPA3), a major component of MCEGs. TNFα in the granular form also shows a higher activity in stimulating endothelial cells as compared to the soluble format. Collectively, these findings reveal MCEGs as functionally active condensates that serve as storage depots while enabling the catalysis of biochemical reactions and mediating immune response.

## RESULTS

### Polyamines form condensates with heparin

Heparin, a glycosaminoglycan, is synthesized by mast cells and stored in mast cell granules^48^, where its concentrations range from a few to a hundred micromolar^67, 68^.

Heparin is a flexible polyanion that, by some estimates, has an excess charge of -3.3 per disaccharide unit^65^. This derives from the numerous sulfate groups that modify the long backbone^69^. Consequently, the heparin interaction with polyvalent counterions is expected to drive the formation of membraneless condensates through complex coacervation^24^. To test this hypothesis, we mixed heparin with mono-, di-, or polyvalent cations at physiologically relevant concentrations. Upon mixing, heparin formed spherical condensates with spermidine and spermine (Fig. 1a), which are polyvalent cations. In contrast, heparin did not form condensates with mono- or divalent cations (Fig. 1a). The tetravalent spermine readily formed condensates with heparin at low concentrations, while the trivalent spermidine required a 5-fold higher concentration to reach the transition threshold (Fig. 1b). We mapped the low-concentration arm of the phase boundary as a function of heparin and spermine concentration. The upper bounds for the mapping were chosen based on known concentrations of heparin^67, 68^ and spermine^70^ in mast cells (Fig. 1c,d and Extended Data Fig. 1a). We also investigated the impact of changing the pH of the buffer and found that this had a minimal effect on condensate assembly in the range of physiologically relevant pH of body fluids (Extended Data Fig. 1b). Besides heparin, chondroitin sulfate is another major glycosaminoglycan synthesized in mast cells^71^, which is structurally close to heparin and also highly negatively charged. We found that chondroitin sulfate formed condensates with spermine, whether alone or mixed with heparin at varying stoichiometries (Extended Data Fig. 1c), suggesting that spermine could form condensates with multiple negatively charged glycosaminoglycans.

**Fig. 1:**
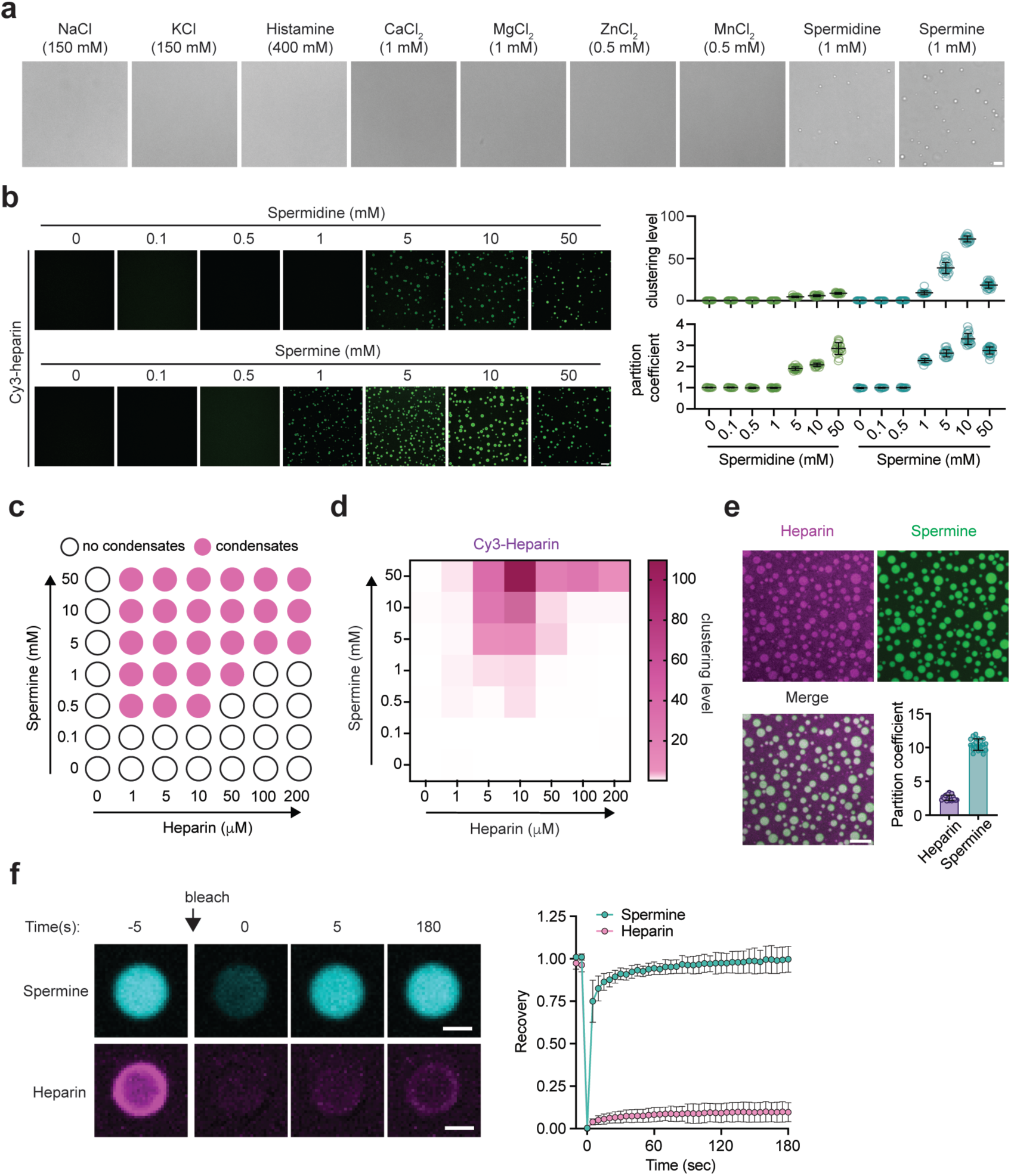
Polyamines form condensates with heparin. **a.** Bright-field images of 5 μM heparin in 50 mM HEPES, pH 7.4 buffer in the presence of cationic salts at the indicated concentrations. These data show that condensates form in the presence of the tri- and tetramines namely spermidine and spermine, respectively. Scale bar, 10 μm. **b.** Condensation of 50 μM FITC-labeled heparin in the presence of increasing concentrations of spermidine and spermine (left panel). The clustering level of heparin was quantified as normalized variance of heparin fluorescence, and the partition coefficient of heparin was quantified as ratio of heparin fluorescence in dense phase against dilute phase. n = 15 individual images/group. Scale bar, 10 μm. **c.** Low concentration arm of the phase boundary mapped in terms of heparin (abscissa) and spermine (ordinate) concentrations. The open circles are concentrations corresponding to the one-phase regime, and solid circles correspond to concentrations that place the system in the two-phase regime defined by coexisting dense and dilute phases. **d.** Quantification of clustering level of heparin condensates as shown in **c** (n = 15 individual images/group). **e.** Fluorescence Images of heparin-spermine condensates formed by 100 μM heparin (added 2 μM Cy5-heparin, magenta) and 10 mM spermine (added 2 μM BODIPY-spermine, green). Partition coefficients (dense phase intensity/dilute phase intensity) of heparin and spermine in condensates were quantified in the lower right histogram, n = 58 individual condensates. Scale bar, 10 μm. **f.** FRAP of heparin-spermine condensates. Right panel showed the time course of fluorescence recovery of Cy3-labeled heparin or BODIPY-spermine in condensates formed by a total of 50 μM heparin and 5 mM spermine. N = 17 (heparin) or 21 (spermine) individual condensates. Scale bar, 2 μm. Data are presented as Mean ± SD. One representative experiment from 3 independent experiments was shown.

Next, we implemented microscopy approaches to further analyze the features of the spermine-heparin condensates. Using Cy5-labelled heparin and BODIPY-labelled spermine, we observed that spermine co-condensed with heparin, attaining an enriched partitioning in the condensed fraction as heparin (Fig. 1e). These data suggest that heparin and spermine form condensates via asymmetrical complex coacervation^24, 27, 29^. The asymmetrical complex coacervates can be formed between smaller, albeit polyvalent cations and larger, high charge-density polyanions, and therefore tend to generate an asymmetry of molecular motions^34, 35, 72, 73^. We used fluorescence recovery after photobleaching (FRAP) to probe for the possibility of asymmetry of molecular motions. We found minimal recovery of the heparin fluorescence following photobleaching of the condensate, in contrast to the efficient recovery of spermine fluorescence (Fig. 1f), suggesting very limited exchange of heparin between the condensed and dilute phase. This contrasts with the efficient recovery of spermine fluorescence, thus validating the expected asymmetry of molecule motions.

### Spermine is enriched in mast cell extracellular granules (MCEGs)

Next, we sought to visualize spermine and examine whether spermine colocalizes with heparin in MCEGs. We incorporated BODIPY-labelled spermine into granules synthesized in bone marrow-derived mast cells (BMMCs)^74^. The successful incorporation of spermine was confirmed by the co-localization of BODIPY-spermine with tryptase and TNFα, two major components of mast cell granules (Extended Data Fig. 2a). These spermine-labeled cells were then activated with anti-TNP IgE and TNP-BSA (IgE/Ag) for 30 mins. The released granules in the medium were captured by dark avidin-coated beads (Fig. 2a). Spermine was detected within captured granules and showed an overlap with heparin (labeled by Sulforhodamine 101-conjugated avidin^57^) (Fig. 2a). Further, flow cytometric analysis revealed that the majority of MCEGs (>95%) contained both spermine and heparin (Extended Data Fig. 2b, upper panels). Their levels remained mostly stable within individual MCEGs even after 4 days when placed at 37°C. Moreover, the MCEG number remained similar within this timeframe (Extended Data Fig. 2c). Together, these data demonstrated that MCEGs are stable entities and that spermine and heparin are core components of MCEGs.

**Fig. 2:**
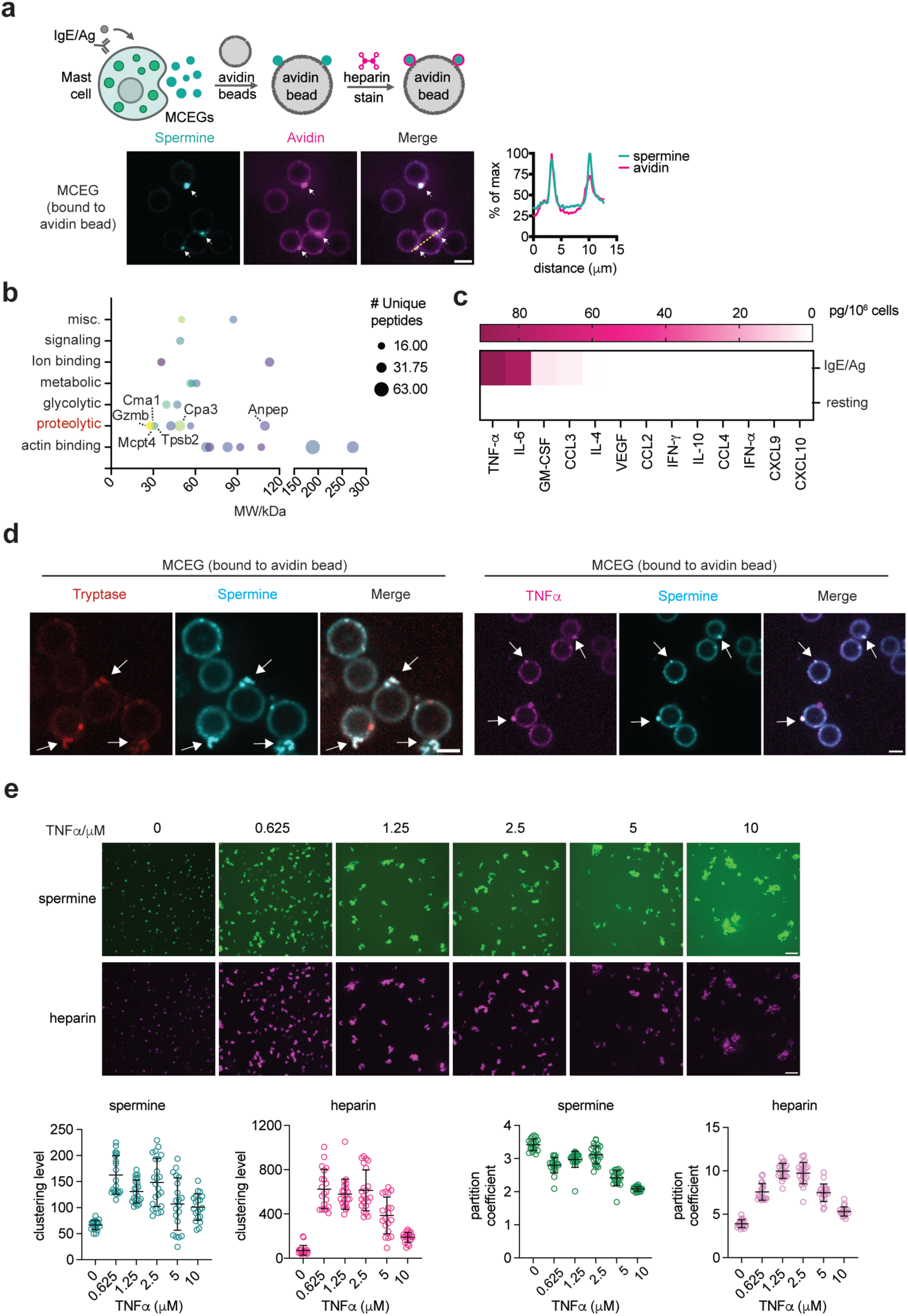
Spermine is enriched in mast cell extracellular granules (MCEGs). **a.** Bone marrow-derived mast cells (BMMCs) loaded with BODIPY-spermine (green) were stimulated with anti-TNP IgE plus TNP-BSA (IgE/Ag) for 30 min. Released granules were captured by avidin beads (schematic workflow in the upper panel) and subjected to Avidin-sulforhodamine 101 (Avidin, magenta) staining and fluorescent imaging (lower left panel). Lower right panel, quantification of fluorescence intensity of BODIPY-spermine and Avidin along the indicated yellow dashed line in the merged image. Scale bar, 5 μm. **b.** Top 25 proteins identified in extracellular granules by mass spectrometry analysis were plotted against their molecular weight. Abbreviations are: Gzmb, GranzymeB; CMA1, chymase; Mcpt4, mast cell protease 4; Cpa3, CPA3; Tpsb2, Tryptase beta-2; Anpep, Aminopeptidase N. **c.** TNFα, IL-6, CCL3, and GM-CSF concentrations in the extracellular granules released by 1 million resting and IgE/Ag stimulated BMMCs were measured by LEGENDplex. N = 2 biological replicates. **d.** MCEGs released from mCherry-Tryptase (red) or mCherry-TNFα (magenta) expressed BMMCs pre-loaded with 20 μM BODIPY-spermine (green) were collected by avidin beads and subjected to confocal imaging. White arrow denotes MCEGs. Scale bar, 5 μm. **e.** Heparin-spermine condensate formation with TNFα. Condensates were formed with 50 μM heparin and 5 mM spermine in the presence of indicated TNFα concentrations. Clustering levels and partitioning coefficients of spermine and heparin were quantified and plotted in lower panels. Data represent analysis of n > 20 individual images per group. Scale bar, 10 μm. Data are presented as Mean ± SD. One representative of three independent experiments was shown.

To determine what protein mediators are retained in MCEGs, we performed mass spectrometry analysis on MCEGs and found that several proteases, including tryptase, chymase, CPA3, and granzyme B, were among the most abundant proteins identified (Fig. 2b and Supplementary Table 1). These proteins are within a subset of proteins identified in previous proteomic studies performed on whole mediators released by mast cells, which include both granular and soluble mediators^75^. Because of the small sizes of cytokines and chemokines, they might not be effectively detected by mass spectrometry. Therefore, we performed a multiplex cytokine profiling assay and found the presence of TNFα, IL-6, GM-CSF, and CCL3 in MCEGs (Fig. 2c).

We further validated that spermine colocalized with both tryptase and TNFα, the two major components within MCEGs (Fig. 2d), and that spermine and heparin were detected in nearly all TNFα- or tryptase-containing MCEGs (Extended Data Fig. 2b, lower panels). Furthermore, FRAP analysis on TNFα in both MCEGs and heparin-spermine condensate revealed minimal fluorescence recovery following photobleaching, indicating that TNFα was stably retained in the granules (Extended Data Fig. 2d-e).

The high stability of MCEGs likely arises from interfacial ionic crosslinks^76^ formed between heparin and spermine, and/or from contributions of protein components. To determine if protein components influence the assembly of heparin-spermine condensates, we included recombinant TNFα, one of the key protein components of MCEG, in the condensation assay and found that TNFα promoted the condensation of both heparin and spermine (Fig. 2e), suggesting the protein components would contribute additional interactions with heparin, spermine, or both that reinforce the stability of MCEGs structure. Notably, TNFα distinctly modulated the partitioning of heparin and spermine into condensates. While spermine partitioning into condensates decreased progressively with increasing TNFα concentration, heparin partitioning was enhanced at intermediate TNFα concentrations (0.625-2.5 μM) (Fig. 2e). This differential partitioning was likely due to competitive binding between TNFα and spermine for heparin, given the presence of positively charged patches on the TNFα surface^77^ .

### Depletion of spermine impedes MCEG assembly

Having observed that spermine is enriched in MCEGs, we sought to assess if spermine is required for MCEG assembly. Spermine is synthesized in the cytosol through the polyamine biosynthesis pathway^78^, and therefore the cellular spermine level can be reduced through a chemical inhibitor N-(3-Amino-propyl)cyclohexylamine (APCHA) against spermine synthase^79^. The reduction of spermine was validated by flow cytometry using an antibody against spermine (Extended Data Fig. 3a). Of note, we selected an intermediate concentration of APCHA to avoid cellular toxicity from the drug at high concentrations.

Next, we determined how depleting spermine affects MCEG assembly. In addition to inhibiting spermine synthesis, we also included reserpine, which inhibits the polyamine transport into granules^80^, and bafilomycin A1, which serves as a positive control that inhibits the vacuolar-type ATPase and disrupts the mediator storage in mast cell granules^81^. We treated BMMCs with the above drugs, followed by the release of intracellular granules to generate MCEGs using anti-TNP IgE and TNP-BSA (IgE/Ag). These are stimuli for Fc epsilon Receptor I (FcɛRI) that are commonly used to release mast cell granules^50, 51^. MCEGs were captured by avidin beads and the number of MCEGs per bead was quantified (Extended Data Fig. 3b). Spermine inhibitors caused a profound reduction in the released granules (Fig. 3b). The total protein amount in MCEGs was also decreased (Fig. 3c). Western blot analysis revealed that the levels of TNFα, IL-1β, CPA3, and tryptase in MCEGs were much lower in granules derived from APCHA- or reserpine-treated cells compared to the vehicle treated cells (Fig. 3d). Importantly, the release of these mediator in soluble form (Fig. 3e) and the expression of these mediators were not affected by spermine inhibitors (Extended Data Fig. 3c), suggesting that these inhibitors primarily affect mediator release in granule form rather than in soluble form.

**Fig. 3:**
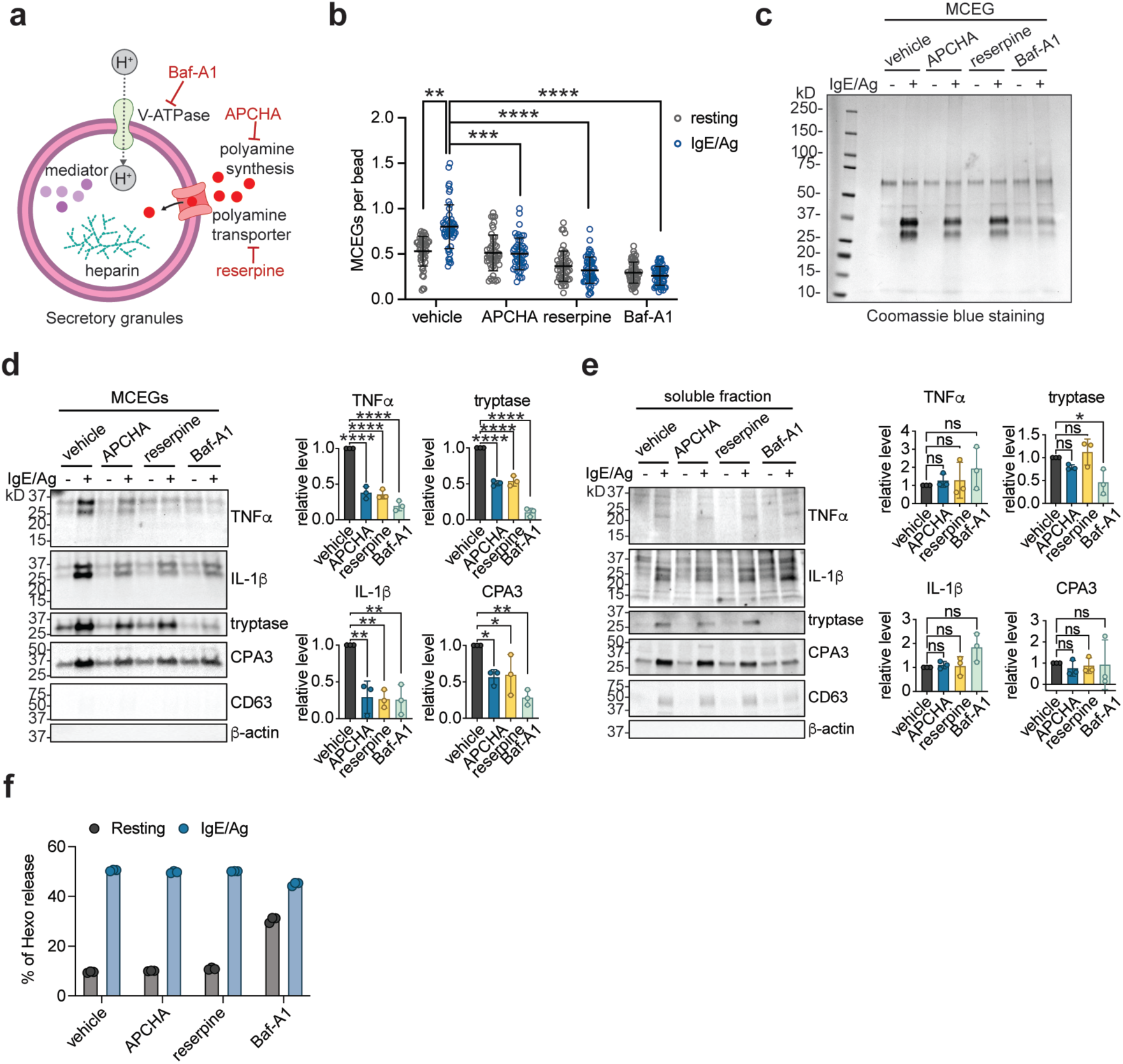
Depletion of spermine impedes MCEG assembly. **a.** Schematic illustration of the chemical inhibitors used in this study and their targets. **b.** MCEGs released from chemical inhibitors-treated BMMCs before and after anti-TNP IgE plus TNP-BSA (IgE/Ag) stimulation were collected with avidin beads. Granules were visualized by Avidin-sulforhodamine 101 staining and the number of granules on each bead was enumerated. N = 50 images per group. **c.** Protein lysates prepared from MCEGs released by 5 million inhibitor-treated BMMCs were analyzed by SDS-PAGE and Coomassie blue staining. **d**-**e** Western blot analysis of mast cell mediators in MCEGs (d) and in the soluble fraction (e). Band intensities for each mediator released in MCEGs (d) and soluble fractions (e) after IgE/Ag stimulation were quantified and normalized to the corresponding mediator in whole cell lysates (Extended Data Fig. 3c). Note that TNFα and IL-1β appeared as multiple bands on the blot because of different isoforms and post-translational modifications^112–115^. These bands were included in quantification. All normalized values were expressed as fold change relative to the vehicle control under IgE/Ag stimulation (set as 1) and plotted in the right panels (each data point represents one independent experiment, n=3). **f.** Degranulation capacity of inhibitors-treated BMMCs was analyzed by the β-hexosaminidase assay. N = 3 technical replicates. Data are presented as Mean ± SD. One representative of two (**b**) or three (**c**-**f)** independent experiments was shown. ns *p* > 0.05, * *p* < 0.05, ** *p* < 0.01, *** *p* < 0.001.**** *p* < 0.0001.

The release of β-hexosaminidase is commonly used as a marker for mast cell degranulation^48^. Of note, the released β-hexosaminidase was primarily found in the soluble supernatant of activated mast cells, but not in MCEGs (Extended Data Fig. 3d), suggesting that β-hexosaminidase was predominantly exocytosed in a soluble form. This provided us with another opportunity to test if spermine specifically affects the presence of mediators in MCEGs or if it affects the secretion of soluble mediators as well. As a result, we found that the β-hexosaminidase levels in the supernatant of APCHA- and reserpine-treated BMMCs were comparable to the vehicle controls, indicating that the extent of mast cell degranulation was not affected by these inhibitors (Fig. 3f). Taken together, using inhibitors against orthogonal protein targets, we demonstrated that spermine selectively regulates the MCEG assembly without affecting the release of soluble mediators.

### Heparin-spermine condensates create a metal ion-rich, alkaline environment for enhanced activity of CPA3

Traditionally viewed as passive mediator storage units, MCEGs house a plethora of mediators with biochemical and signaling activities^48^. To understand the functional consequences of granule assembly, we first asked whether the biochemical activities of mediators are influenced by granule assembly. We focused on CPA3, a carboxypeptidase highly enriched in MCEGs (Fig. 3d), and compared its enzymatic activity between the soluble and granule formats.

BMMCs were stimulated by anti-TNP IgE and TNP-BSA, and the released granules were harvested and examined for CPA3 activity using a CPA3-specific substrate M-2245. To obtain the soluble form of CPA3 from MCEGs, the harvested granules were treated with heparinase I, which breaks down heparin^82^. Western blot analysis showed that after heparinase I treatment, CPA3 was almost completely released into the soluble supernatant (Fig. 4a). Interestingly, CPA3 in this soluble format showed reduced enzymatic activity as compared to that in the intact granule (Fig. 4a). We verified that the presence of heparinase I did not affect the readout of this assay (Extended Data Fig. 4a). Therefore, these data suggested that CAP3 in the soluble format showed a lower protease activity than in granules.

**Fig. 4:**
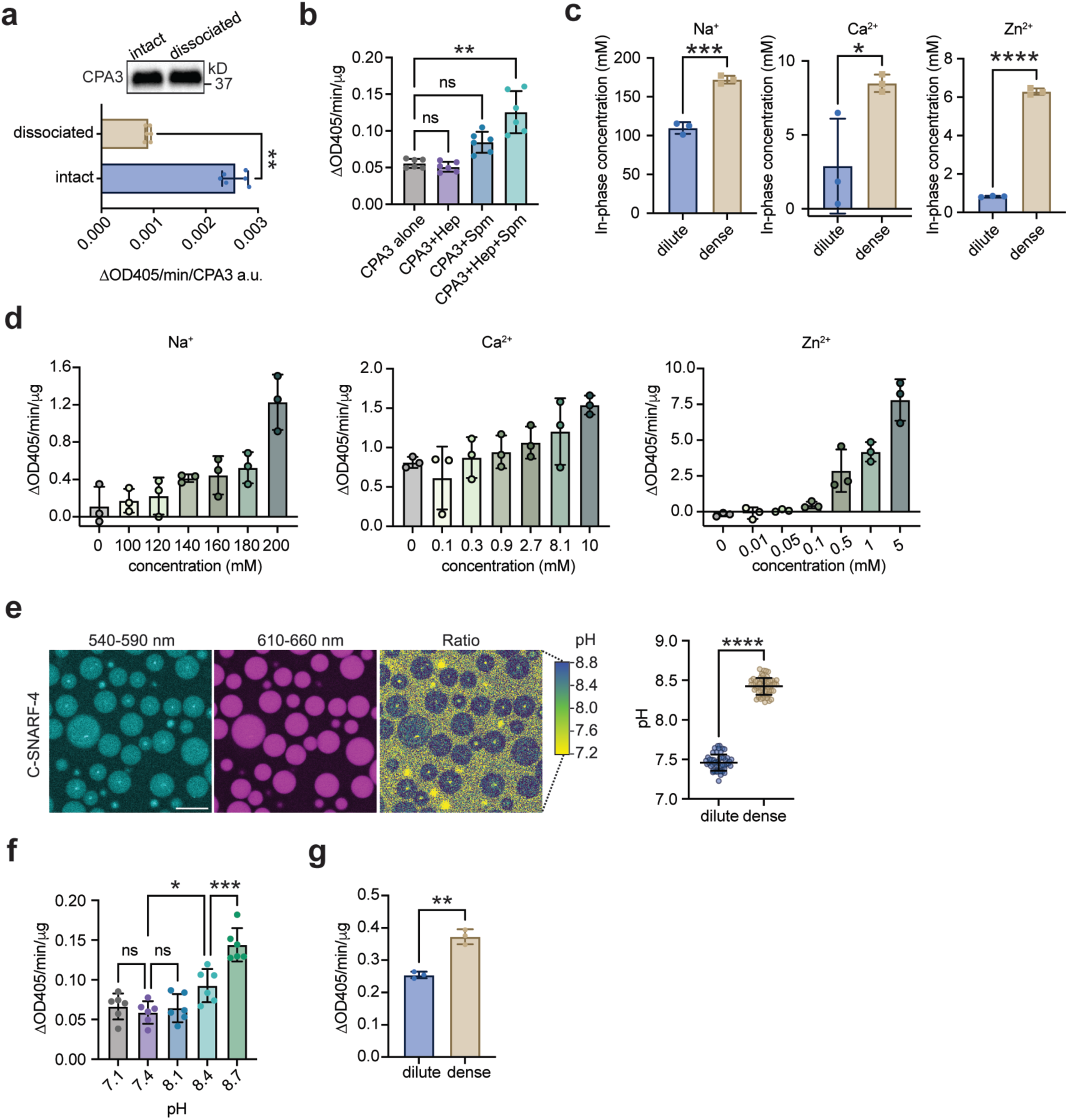
Heparin-spermine condensates create a metal ion-rich, alkaline environment for enhanced activity of CPA3 **a.** CPA3 protease activities of intact native granules and of dissociated granules (treated with heparinase I) were measured using chromogenic substrate and normalized to CPA3 densitometry unit from western blot analysis. N = 6 biological replicates. **b.** Protease activities of recombinant CPA3 in either tyrode’s buffer, combined with heparin, spermine, or in heparin-spermine condensates were measured. N = 6 biological replicates. **c.** ICP-MS analysis of the ion concentrations in the dilute phase and the dense phase of heparin-spermine condensates. N = 3 biological replicates, each with 3 technical replicates. **d.** Protease activities of recombinant CPA3 in 50 mM HEPES, pH 7.4 buffer containing varying concentrations of NaCl, CaCl_2_, or ZnCl_2_ as indicated. N = 3 biological replicates. **e.** pH difference between the dense phase and surrounding dilute phase of heparin-spermine condensates was determined with ratiometric C-SNARF-4 dye. pH values were calculated from the ratio of fluorescence intensities at 540-590 nm and 610-660 nm using a calibration curve (n = 59 condensates, right panel). Scale bar, 10 μm. **f.** Protease activities of recombinant CPA3 at different pH. N = 6 biological replicates, one representative from two individual experiments was shown. **g.** Protease activities of recombinant CPA3 in pH- and salt-adjusted buffers mimicking the dilute or dense conditions of heparin-spermine condensates. N = 3 biological replicates Data are presented as Mean ± SD. One representative of two (**a**-**b**, **e**) or three (**d**, **f**-**g**) independent experiments was shown. ns *p* > 0.05, * *p* < 0.05, ** *p* < 0.01, *** *p* < 0.001, **** *p* < 0.0001.

To address the concern that the reduced protease activity of soluble CPA3 could be due to a loss of interaction or regulation by other proteins in the granules, we performed a similar comparison using reconstituted granules in which we could accurately control the compositions. Recombinant CPA3 was mixed with heparin alone, spermine alone, or heparin plus spermine which reconstitutes the granule condensates. CPA3 was confirmed to be recruited to the reconstituted condensates (Extended Data Fig. 4b). We found that the protease activity of CPA3 in granules was higher than CPA3 either on its own or combined with heparin or spermine, where no condensate formation was observed (Fig. 4b).

The enhanced protease activity of CPA3 in granules could be due to a high local concentration of enzyme or a distinct chemical environment in granules. We determined the protease activity of CPA3 with increasing concentrations (up to 50-fold). However, we found a dose-dependent decrease, rather than an increase, in its specific activity (activity per molecule) (Extended Data Fig. 4c), disfavoring a concentration-dependent mechanism explaining the activity increase.

On the other hand, previous reports showed that the protease activity of CPA3 is regulated by pH and cations^83^. Recent studies suggested condensates that form both in vitro and in cells are characterized by interphase pH and ion gradients, thus giving rise to distinct electrochemical environments within condensates that are system-specific and age-dependent^31, 84–87^. Motivated by these observations, we asked if the heparin-spermine condensates create an electrochemical environment that is distinct from the surrounding dilute phase. We reconstituted the heparin-spermine condensates in a salt buffer comprising metal ions (Na^+^, Ca^2+^, Zn^2+^) that were commonly found in the extracellular space. The dilute and dense phase were separated from the bulk solutions by centrifugation. The cation abundance was measured using inductively coupled plasma mass spectrometry (ICP-MS)^85, 88^. We found that the concentrations of Na^+^, Ca^2+^, and Zn^2+^, were higher in the dense phase, showing increases of ∼1.5-fold for Na^+^, ∼3-fold for Ca^2+^, and ∼7-fold for Zn^2+^ when compared to the dilute phase (Fig. 4c). Note that the buffer contained physiologically relevant concentrations of Na^+^ and Ca^2+89, 90^ but higher than normal Zn^2+^ concentration because the physiological relevant concentration of Zn^2+^ (nM to μM range^91^) is below the detection limit of ICP-MS. Further, we demonstrated that the enzymatic activities CPA3 were enhanced with increased metal ion concentrations (Fig. 4d). These data suggested that the enrichment of metal ions in the heparin-spermine condensates increases CPA3 activities.

In parallel, we determined the pH within heparin-spermine condensates using a ratiometric sensor C-SNARF-4^31, 86, 92^. The internal pH is another key parameter that defines the electrochemical environment of condensates^31, 86^. We found that pH inside the heparin-spermine condensate was one unit higher than the outside (pH 8.4 versus 7.4) (Fig. 4e), implying a 10-fold lower concentration of protons within condensates. Notably, an alkaline pH favored enhanced CPA3 activity (Fig. 4f). Using buffers mimicking both the pH and metal ion concentrations observed in dense and dilute phase samples, we found that CPA3 in the dense phase buffer exhibited higher activity than in the dilute phase buffer (Fig. 4g). Together, these results revealed a unique electrochemical environment inside heparin-spermine condensate, which regulate the biochemical activities of mediators.

### Condensation reverses the inhibition by heparin or spermine on TNFα-triggered endothelial activation

MCEGs enter the lymphatic and blood vessels after being released from activated mast cells in tissues^41, 60^. Given that they contain mediators such as cytokines and chemokines, circulating MCEGs represent unique forms of messengers, packaged in a distinct electrochemical environment, that convey an immune signal remotely without trafficking of cells. We asked whether the granular delivery of a mediator would have a distinct functional outcome for cell-to-cell communication as compared to its soluble format. To employ an experimental system wherein we can accurately control the condensation level and composition, we reconstituted synthetic granules in vitro with heparin, spermine, and TNFα, a potent immune modulator identified in mast cell granules^41, 60, 93, 94^. The successful loading of TNFα onto the heparin-spermine condensates was confirmed by confocal fluorescence microscopy (Fig. 5a). As a control, mixing TNFα with either heparin or spermine (soluble TNFα) did not induce condensate formation of TNFα. Equal amounts of TNFα in condensates or in soluble formats were added to human umbilical vein endothelial cells (HUVECs) (Fig. 5b). Substantial numbers of TNFα puncta were observed on HUVECs treated with TNFα in condensates, but not with soluble TNFα. Flow cytometric analysis revealed prolonged retention of condensed TNFα on HUVEC cells, lasting from 1 hr to 24 hr. In contrast, the retention of soluble TNFα was considerably lower (Fig. 5c). Interestingly, the combination of TNFα and spermine also increased TNFα retention on HUVEC cells after 24 hr, although to a lesser extent than condensed TNFα. This effect may be attributed to the subsequent interaction of TNFα and spermine with heparan sulfate on the HUVEC surface, thereby promoting its retention.

**Fig. 5:**
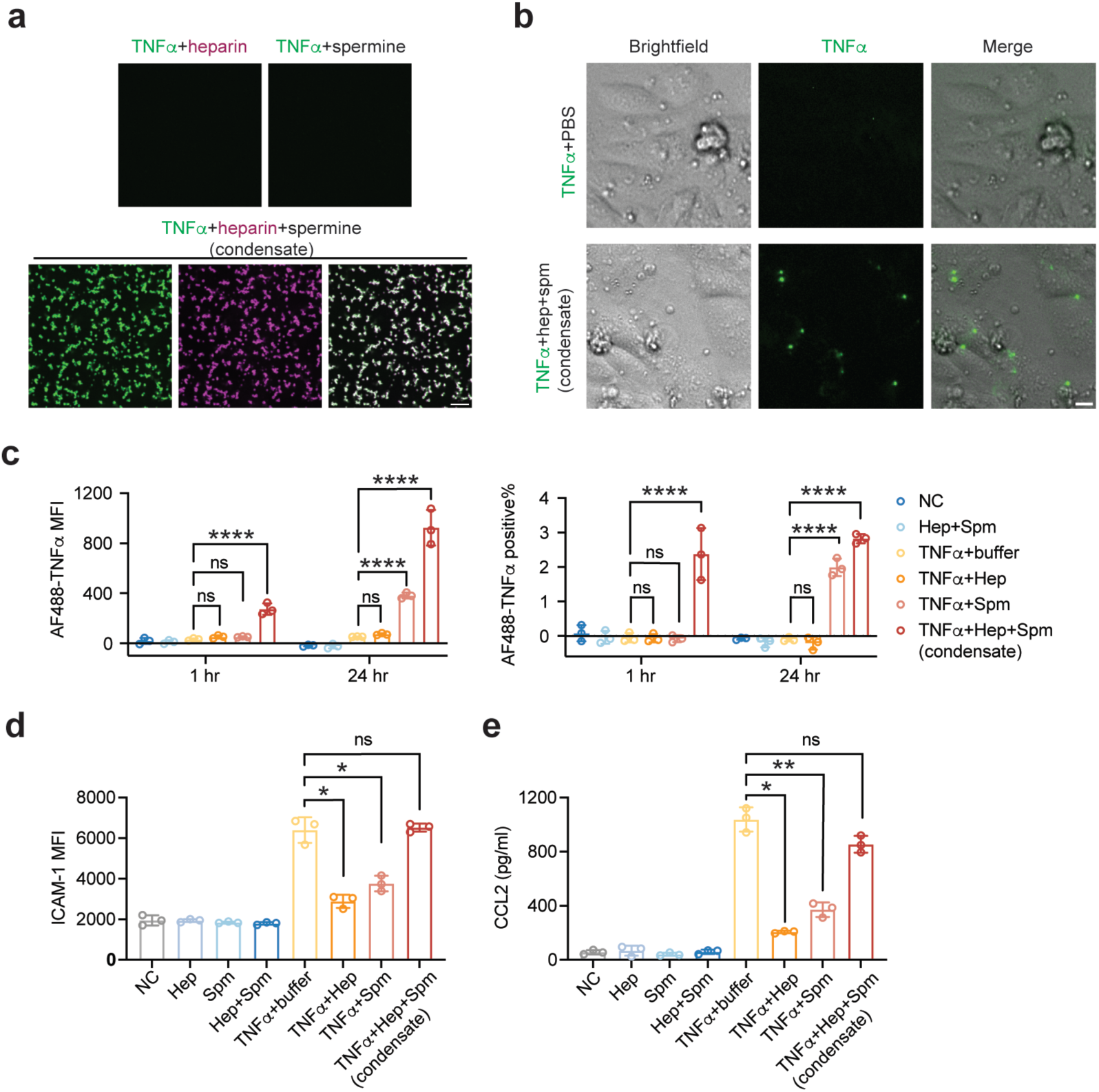
Condensation reverses the inhibition by heparin or spermine on TNFα-triggered endothelial activation. a. Fluorescence images of 100 nM AF488 labeled-TNFα (green) in the presence of 50 μM heparin (premixed with 2 μM Cy5 labeled-heparin, magenta), 1 mM spermine, or in heparin-spermine condensates. Scale bar, 10 μm. **b.** Bright-field and fluorescence Images of HUVEC cells after 24 hr treatment with AF488 labeled-TNFα (1 ng, green) in PBS buffer or in heparin-spermine condensates. Hep, heparin; Spm, spermine. Scale bar, 10 μm. **c**-**e**. HUVEC cells were treated with 1 ng AF488 labeled-TNFα in either PBS buffer, combined with heparin, spermine, or in heparin-spermine condensates. **c**, Mean fluorescence intensity of AF488 labeled-TNFα on HUVEC cells, as well as the percentage of AF488 labeled-TNFα-positive cells after 1 hr and 24 hr were analyzed by flow cytometry. **d**, Surface ICAM-1 expression on HUVEC cells was analyzed by flow cytometry. **e**, CCL2 levels in the cell supernatant were measured by ELISA. N = 3 technical replicates. Data are presented as Mean ± SD. One representative of three independent experiments was shown. ns *p* > 0.05, * *p* < 0.05, ** *p* < 0.01, **** *p* < 0.0001.

Next, we examined TNFα-induced HUVEC activation. Consistent with previous reports^95^, heparin inhibited the TNFα-induced ICAM-1 upregulation and CCL2 production (Fig. 5d-e). Spermine also elicited similar inhibition on TNFα-induced activation. In contrast, condensed TNFα, which contains both heparin and spermine, induced comparable ICAM-1 surface expression and CCL2 production as soluble TNFα in PBS buffer. This suggests that condensed TNFα is able to reverse the inhibition of heparin and spermine on the functions of TNFα.

## Discussion

This work shows that mast cell extracellular granules (MCEGs) form by co-condensation of polyamines with glycosaminoglycans (Fig. 6). First, heparin—used as a model glycosaminoglycan—readily forms co-condensates with polyamines *in vitro*. Second, depleting cellular polyamines reduces MCEG production. Third, polyamines are not only imported into the intracellular granules that generate MCEGs but are also present within the extracellular MCEGs, consistent with a co-condensation mechanism. This discovery opens new avenues for applying the principles of biomolecular condensation to biological processes beyond the intracellular space, particularly for membraneless bodies that mediate cell-to-cell communication and immune responses. Further, MCEGs present a powerful model for understanding how the internal electrochemical environment of a condensate dictates the activity and function of its cargo proteins.

**Fig. 6:**
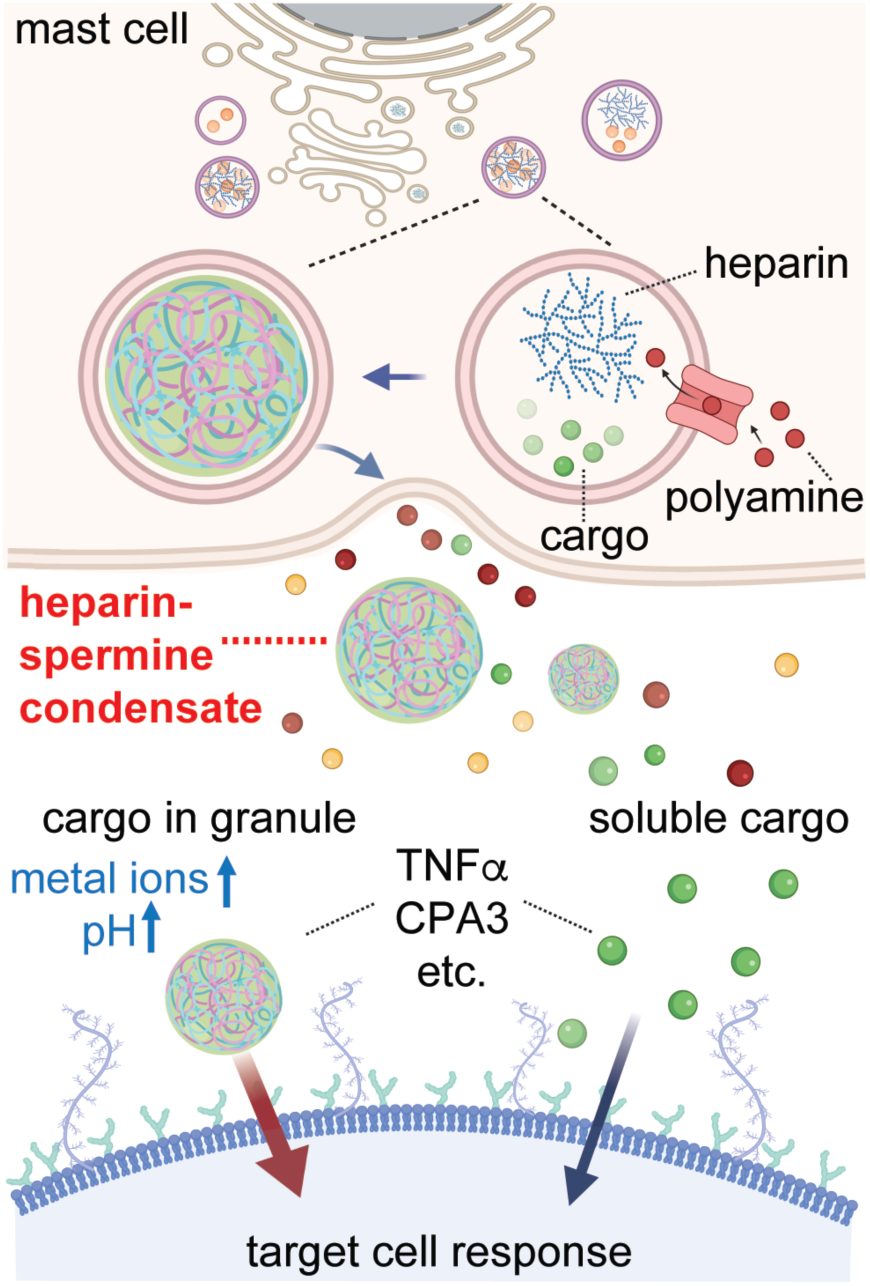
Proposed mechanism and function of MCEGs. MCEGs are assembled through electrostatic interactions between heparin and spermine. The storage of proteases and cytokines within MCEGs relies on heparin-spermine condensate. Reconstituted granules comprising heparin and spermine can efficiently capture mediators such as CPA3 and TNFα, enhancing their respective enzymatic and bioactive potency. The enhanced functionality stems from the distinct electrochemical microenvironment formed within the heparin-spermine condensate, marked by elevated pH and enriched metal ions. Collectively, MCEGs represent functionally active condensates that serve as storage depots while facilitating biochemical reactions and mediating immune responses.

To definitively address MCEG function, we established a reconstitution system that recapitulates their assembly, cargo retention, and chemical environment. Using this system, we demonstrated that both native MCEGs and reconstituted CPA3 granules exhibit significantly higher protease activity than their dissociated or soluble counterparts. We characterized the distinct electrochemical properties of these spermine-heparin condensates. Consistent with the asymmetric ion partitioning observed in protein and RNA condensates^85, 87, 92^, the spermine-heparin condensates exhibit an alkaline pH and a high cation concentration that directly promote enhanced CPA3 activity.

MCEGs have traditionally been viewed as storage depots for fully active pre-formed mediators such as TNFα during long-distance cell-to-cell communication^41, 60^. The thinking has been that the retention of mast cell mediators, such as CPA3, tryptase, and chymase from MCEG is contingent on their basic surface patches that underly the electrostatic interactions with the negatively charged proteoglycan core^96–104^. Our work, which revealed a unique electrochemical milieu within MCEGs, suggests that MCEGs function as specialized biochemical reaction hubs. This finding demonstrates that the functional identity of MCEGs is defined by their unique internal chemical environment. Rather than being simple storage depots, they are specialized hubs where the electrochemical conditions created by complex coacervation directly regulate mediator activity. We have thus uncovered a form of extracellular signaling where the message is not just the molecule, but the specific physicochemical context in which it is delivered. This capacity to create potent, self-contained signaling hubs outside the cell represents a fundamental advance in our understanding of the function of biomolecular condensates in cell-to-cell communication.

This notion was strengthened by our finding that granular delivery of cargo protein TNFα elicits prolonged retention and sustained response on target cells, unlike soluble TNFα. The functional differences between granular and soluble forms imply the contribution from structural components heparin or spermine to its cargo functions. Previous studies using cell-free systems showed that heparin association enhances the functions of its binding partners. This is achieved through various mechanisms, including direct modulation of oligomerization status for activation (e.g., chemokines CCL3 and CCL5^105^), protection against proteolytic degradation (e.g., tryptase ^106^), or bringing substrate into close proximity for cleavage (e.g., chymase^107^). Conversely, heparin inhibits the functions of cytokines, such as IL-6^108^, IFNγ ^109^, IL-10^110^, and TNFα ^95^, by either masking their conjugations with corresponding receptors or by inducing negative effector proteins for downstream cytokine signaling. Our findings corroborated the inhibitory effect of heparin on soluble TNFα^95^. Notably, the presence of spermine to form TNFα-heparin-spermine condensates reversed the heparin inhibition, highlighting the functional interplay between the heparin/spermine matrix and cargo proteins within a granule. The functional significance of such a matrix environment is further supported by studies using synthetic heparin-chitosan particles, where TNFα in synthetic particles exhibits sustained release kinetics^111^. These findings establish that the coacervate matrix of MCEGs dictates the ultimate signaling output of its cargo, enabling a form of context-dependent signaling. Therefore, the unique functionality conferred to mast cells by MCEGs arises from their capacity to actively tune mediator activity, transforming both the kinetics and potency of immune messages.

## Methods Mice

All Animal care and experiments were approved by the Yale University Animal Care and Use Committee and in accordance with the US National Institutes of Health guidelines. Mice were housed at the Yale University animal facilities in specific pathogen-free conditions, maintained on a 12-h light-dark cycle in a temperature (22°C)- and humidity-controlled room. Female C57BL/6 and BALB/c mice were obtained from Envigo and used at 6–12 weeks of age for all experiments.

### Cell culture

To generate bone marrow-derived mast cells (BMMCs), bone marrow cells were flushed from mouse femurs and tibias and cultured in RPMI-1640 medium containing 20% FBS (Biowest), 1x Penicillin-Streptomycin-Glutamine, HEPES, non-essential amino acid, 2-mercaptoethanol (Gibco), 20 ng/mL recombinant murine stem cell factor, and 20 ng/mL recombinant murine IL-3 (Sino Biologicals). Cells were seeded at a density of 1× 10⁶ cells/mL and maintained at 37°C in a 5% CO_2_ humidified incubator. After 4-6 weeks, >95% of cells were confirmed mature by CD117 and FcεR1a expressions with flow cytometry. HEK293T cells were maintained in DMEM medium supplemented with 10% FBS and 1 x Penicillin-Streptomycin-Glutamine in a 5% CO_2_ humidified incubator at 37°C. Human umbilical vein endothelial cells (HUVEC; ATCC) were cultured in Vascular Cell Basal Medium supplemented with Endothelial Cell Growth Kit-BBE (ATCC) and incubated at 37°C with 5% CO_2_. Inhibitors including N-(3-Aminopropyl)cyclohexylamine^79^ (APCHA, 0.5 mM, TCI chemicals), reserpine^80^ (10 μM, Cayman Chemical), or bafilomycin A1^81^ (Baf-A1, 20 nM, Cayman Chemical), were added to mast cell culture three days before subsequent experiments.

### Plasmid construction and lentiviral transduction

DNA fragments encoding mouse Mcpt6 or Tnfa were synthesized (Genescript) and cloned into a pHR lentiviral vector in-frame with a mCherry tag at C-terminus under an SFFV promoter. For piggyBac transposon-based system, DNA fragments encoding proteins of interest were cloned into the PB-T-PAF vector as previously described^116^. Sequences of all plasmids were verified by Sanger sequencing. Lentivirus was generated by transfection of HEK 293T cells with packaging plasmids pMD2.G and psPAX2 (Addgene plasmid #12259 and #12260) using linear polyethyleneimine hydrochloride (PEI) MAX, MW. 40,000 (Polysciences). After 48 hr, viral supernatants were collected, centrifuged, filtered using 0.45-μm filters, and concentrated using Lenti-X concentrator (Takara) following the manufacturer’s instructions. Virus pellets were resuspended in PBS, aliquoted and stored at −80°C until use. BMMCs were transduced with lentivirus by spinoculation at 1200 ×g for 90 min at 32 °C. Cells were then maintained in fresh complete medium for at least a week before use.

### Mast cell degranulation

BODIPY conjugated-spermine (20 µM, Merck, BODIPY FL conjugate) was added to BMMCs cell culture for two days. The BODIPY-spermine-loaded BMMCs were then stimulated with 1 µg/mL anti-TNP IgE (BioLegend, clone TNP7) plus 100 ng/mL TNP-BSA (Cayman Chemical) (IgE/Ag) for 30 min at 37°C in Tyrode’s buffer (10 mM HEPES, 130 mM NaCl, 5 mM KCl, 1.4 mM CaCl₂, 1 mM MgCl₂, 5.6 mM glucose, pH 7.4) containing 10 µg/mL Avidin-sulforhodamine 101 (Avidin-SR101; Sigma-Aldrich).

Post-stimulation, cell supernatants were centrifuged at 500 × g for 5 min to remove cellular debris. Released heparin-containing granules from IgE/Ag-activated BMMCs were captured using avidin polystyrene beads (6.0-8.0 µm, Spherotech), incubated in the same cell supernatant for 30 min at RT on a rocking platform, and then subjected to confocal imaging.

To identify proteins in the MCEGs from IgE/Ag BMMCs, MCEGs were collected by centrifugation at 20,000 × g for 60 min and lysed in NuPAGE LDS sample buffer (Thermo Scientific) supplemented with protease inhibitors. After boiling at 100°C for 5 min, granule lysates were loaded onto a 4-20% gradient gel and run for 3 min until all the samples entered the gel and stacked into a thin band. The gel was then stained with GelCode Blue (Thermo Scientific) according to the manufacturer’s instructions. Protein bands with visible blue staining were excised and subjected to Mass Spectrometry analysis using Q-Exactive Hybrid Quadrupole-Orbitrap Mass Spectrometer (LC-MS/MS) at the Protein Facility of Iowa State University.

### β-hexosaminidase release assay

β-Hexosaminidase activities in the cell fractions were quantified with p-nitrophenyl-N-acetyl-β-D-glucosaminide (NAG, Sigma) as substrate. Cell supernatant, cell lysate, and extracellular granules dissolved in 0.5% Triton buffer were incubated with NAG substrate solution (3.4 mg/mL in buffer containing 17.6 mM Na₂HPO₄, 11.2 mM sodium citrate, pH 4.5) at 37°C for 1 hr. Reactions were stopped by adding glycine buffer (2 M Na₂CO₃, 1.1 M glycine, pH 10). The absorbance was read at 405 nm using a SpectraMax M5 microplate reader (Molecular Devices). Background absorbance was subtracted. A standard curve was prepared using beta-N-Acetyl-Hexosaminidase (NEB, 0–1.6 U/mL) in parallel with the samples. Enzyme activity was calculated from the standard curve.

### Protein expression and purification

HEK 293T cells stably expressing His-tagged TNFα proteins were generated using piggyBac transposon-based expression system following the established protocol^116^. To induce protein expression, 1 μg/mL doxycycline was added to the cell cultures maintained in serum-free DMEM medium for 24 hr. Cell supernatant was then collected and clarified by centrifugation and filtration through 0.45-μm filters. His-tagged TNFα was then bound with Ni-NTA resin (GE Healthcare), washed with buffer containing 20 mM imidazole (50 mM HEPES, pH 7.4, 150 mM NaCl, 1 mM TCEP, 10% glycerol), and eluted with 500 mM imidazole buffer containing 50 mM HEPES, pH 7.4, 150 mM NaCl, 1 mM TCEP, and 10% glycerol. The proteins were subjected to gel filtration chromatography using a Superdex 200 10/300 GL column (GE Healthcare) in equilibrium buffer (50 mM HEPES, pH 7.4, 150 mM NaCl, 1 mM TCEP, and 10% glycerol). Monomer fractions were pooled, labelled with Alexa Fluor 488-maleimide (Lumiprobe) for 2 hr at room temperature (RT), and run through a PD MidiTrap G-25 (GE Healthcare to remove excess dye. Protein purity was verified by SDS-PAGE and Coomassie staining.

### Flow cytometry

Cell surface staining was performed by incubation of cells with antibodies in PBS containing 2% BSA for 30 min at 4°C. The following antibodies were used: PE Cy7-FcχR1α (Clone MAR-1, Biolegend), Alexa Fluor 488-CD117 (Clone 2B8, Biolegend), Alexa Fluor 647-ICAM-1 (Clone HA58, Biolegend). Isotype controls were used to establish positive gates. For intracellular spermine staining, cells were fixed with 4% paraformaldehyde and then permeabilized with 0.5% saponin for 20 min. Cells were rinsed with PBS and incubated with anti-spermine antibody (Abcam, 1:500 dilution in PBS containing 2% BSA) overnight at 4°C, followed by incubation with Alexa Fluor 647 anti-rabbit secondary antibody (1:1000 dilution in PBS containing 2% BSA) for 1 h at room temperature. Cells and MCEGs were analyzed on a 4-laser Cytek Aurora cytometer (Cytek Biosciences). For MCEGs quantification, CountBright Absolute Counting Beads (Invitrogen) were spiked into MCEGs samples at Day 0. MCEGs were acquired under low-speed mode with a fixed volume, then normalized to the counting bead number. Data were processed using FlowJo v10.10.0 (BD Life Sciences).

### *In vitro* condensation assay

Heparin-spermine condensate was formed by mixing spermine and heparin solution in 50mM HEPES, pH 7.4 buffer. To visualize the condensates under fluorescent imaging, BODIPY-spermine, or Cy5-, FITC-, or Cy3-labelled heparin was added to the dark spermine or heparin buffer before mixing, reaching a final concentration of 2 μM. Condensates were incubated for 15 min at 25°C before subsequent experiments. To separate dense phase and dilute phase, bulk condensate sample was subjected to centrifugation at 500 × g for 10 min at RT. The viscous dense phase sample was collected with a positive displacement pipette.

### SDS-PAGE and Western blot analysis

To analyze proteins in the MCEGs and the soluble fraction following IgE/Ag stimulation, cell suspensions were centrifuged at 500 × g for 5 min to remove cells and debris. MCEGs were isolated from the supernatant by centrifugation at 20,000 × g for 60 min. The remaining soluble fraction was concentrated using a 3 kDa molecular weight cut-off ultracentrifugal filter (Amicon). Cell pellets, MCEGs, and soluble fraction were dissolved in NuPAGE LDS sample buffer (Thermo Scientific) supplemented with protease inhibitors, and boiled at 100 °C for 5 min. Proteins were separated on 4–20% Mini-PROTEAN TGX stain-free protein gels (Bio-Rad) by SDS-PAGE. For Coomassie staining, gels were stained with Gelcode blue safe protein stain (Thermo Scientific) for 1 hr, and destained in MilliQ water (Millipore). For western blotting, proteins were transferred onto PVDF membranes (0.22 μm, Millipore), followed by blocking with 5% BSA in PBS supplemented with 0.1% Tween-20 (PBS-T). Membranes were then incubated with primary antibodies (1:1000 dilution) in PBS-T buffer at 4 °C overnight, followed by incubation in HRP-conjugated secondary antibodies (1:10000 dilution) in PBS-T buffer for 1 hr. Protein bands were visualized using SuperSignal ECL substrate (Thermo Scientific) under a ChemiDoc XRS+ imaging system (Bio-Rad). The primary antibodies used were anti-TNFα (clone MP6-XT22, Biolegend), anti-IL-1β (clone 3A6, Cell Signaling Technology), anti-CPA3 (Proteintech), anti-tryptase (clone 3G3, Bioss), anti-CD63 (clone NVG-2, Biolegend), and anti-β-actin (clone 13E5, Cell Signaling Technology). Secondary antibodies used were HRP-conjugated goat anti-rat IgG (H+L), goat anti-rabbit IgG (H+L), and goat anti-mouse IgG (H+L) (all from Thermo Scientific).

### Confocal imaging and FRAP

Confocal imaging and Fluorescence recovery after photobleaching (FRAP) experiments were performed on microscope system composed of a Nikon Ti2-E inverted motorized microscope stand, motorized stage with stagetop Piezo, Lapps system with XY miniscanner for 405 nm FRAP, CSU-X1 spinning disk confocal, Agilent laser combiner with four lines, 405, 488, 561, and 640 nm, and scientific CMOS camera Photometrics Prime 95B. Images were acquired using a Nikon 60× Plan Apo 1.40 NA oil immersion objective with NIS Elements software and analyzed in Fiji (Image J2, 2.16.0/1.54p). Imaging of condensates was performed on mPEG-silane (Laysan Bio) passivated coverslips^117^, which have been pre-cleaned with 5% Hellmanex III (Sigma) and etched with 5 M NaOH. The fluorescence clustering level was quantified using normalized variance (0−²/μ), calculated by dividing the square of the standard deviation by the mean fluorescence intensity across the entire image. The partitioning coefficient, reflecting the enrichment of fluorescence in condensates, was determined by measuring the ratio of fluorescent intensity in dense phases to that in dilute phases. For confocal imaging of BMMCs or MCEGs-bound avidin beads, samples were imaged on a glass-bottom 384-well plate (CellVis). For FRAP, a region of interest (ROI) was photobleached using a 405nm FRAP laser. Fluorescence recovery was monitored over 3 min at 5-sec intervals. Prior to photobleaching, 4 baseline images at 5-sec intervals were recorded. For each time point, the fluorescence of an unbleached region was measured as a reference to correct for nonspecific photobleaching. The fluorescent intensity of bleached ROI over time was quantified with background fluorescence subtracted, corrected by the unbleached region at the same time point, and then normalized to the prebleached intensity of the ROI (set as 1).

### ICP-MS

A bulk condensate sample was prepared by mixing 2 mL of 1 mM heparin solution with 18 mL of 10 mM spermine solution in 50mM HEPES, pH 7.4 buffer supplemented with salts. The final salt concentrations in the mixture were 150 mM NaCl, 1 mM CaCl_2_, 5 mM KCl, 5 mM MgCl_2_, 500 μM ZnCl_2_, and 500 μM MnCl_2_. The bulk sample was centrifuged at 500 × *g* for 10 min to separate the dilute and dense phases. Eight microliters of each phase were pipetted into borosilicate glass tubes containing 0.5 mL nitric acid (70%, trace metal grade, Fisher Chemical). Samples were boiled at 90 °C for 2 hr with glass coverslips placed on the tubes to minimize evaporation, then cooled to RT. Samples were further diluted with 16.5 ml of 2% nitric acid (v/v in deionized water) and analyzed using a NexION 2000 inductively coupled plasma mass spectrometer (PerkinElmer, Waltham, MA, USA) equipped with a dynamic reaction cell (DRC) and autosampler (PerkinElmer), as previously described^85^. For calibration, standard solutions containing a mixture of sodium, potassium, calcium, zinc, magnesium, and manganese at serial concentrations were prepared in 2% nitric acid (v/v). Standards were analyzed before and after dilute/dense phase sample measurements to ensure instrument stability throughout the experiment in addition to a Sc internal standard. Raw counts per second (cps) data for both standards and dilute/dense phase samples were recorded. Data were processed by normalizing cps values to the calibration curves, internal standard, and accounting for dilution factors.

### pH measurement with C-SNARF-4

The ratiometric pH-sensitive dye C-SNARF-4 (Thermo Scientific) was used to measure pH inside and outside condensates, following an established protocol^92^. Samples containing C-SNARF-4 (20 µM) were loaded in a glass-bottom 384-well plate and imaged on a Leica SP8 equipped with an HC PL APO 40x/1.30 oil immersion objective. Imaging was performed using a constant 75% white light laser (WLL) power with excitation at 531 nm. Emission signals were acquired in two channels, 540-590 nm and 610-660 nm. To establish a calibration curve, a series of 50 mM Tris buffers (pH range 7-9) was prepared and mixed with C-SNARF-4 probe (20 µM). Fluorescence images of the calibration samples were acquired under identical imaging conditions. The ratio of fluorescence intensity (540-590 nm/610-660 nm) was calculated in FIJI (ImageJ) and plotted against the actual pH values measured with a calibrated pH meter (Mettler Toledo). Sample pH values were determined by applying the measured fluorescence intensity ratio to the calibration curve.

### CPA3 protease activity assay

CPA3 protease activity was measured using the chromogenic substrate N-(4-Methoxyphenylazoformyl)-Phe-OH potassium salt (M-2245, Bachem). Reactions were initiated by mixing 5 μl of sample (100 nM) with 50 μl of M-2245 substrate stock (1 mM). Absorbance at 405 nm was recorded over 10 min at 15-second intervals using a SpectraMax microplate reader (Molecular Devices). Cleavage of M-2245 results in a decrease in absorbance at 405 nm. The enzyme activity was determined from the initial linear phase of the absorbance-versus-time curve, typically within the first 2 mins of the assay. CPA3-specific activity was expressed as differences in absorbance at 405 nm per minute per μg of CPA3.

## Statistics

Statistical analysis was performed using GraphPad Prism v10.4.0. Data are presented as Mean or Mean ± SD as indicated. When comparing the means of two groups, a two-tailed unpaired Student’s t-test or Mann–Whitney U test for nonparametric data was used. When comparing more than two groups, a one-way or two-way ANOVA followed by multiple comparison test was used. \**p* < 0.05, ** *p* < 0.01, *** *p* < 0.001, **** *p* < 0.0001, ns *p* > 0.05.

## Acknowledgements

We thank Drs. S. Galli, R. Sagi-Eisenberg, S. Abraham, Y. Dai for feedback on the manuscript. X.S. was supported by an American Cancer Society Research Scholar Grant 135926, the NIGMS MIRA program R35 GM138299, the Gabrielle’s Angel Foundation Medical Research Award, the Pershing Square Sohn Prize for Young Investigators in Cancer research, the NCI Exploratory/Developmental Research Grant R21 CA286364, the NIH Exploratory/Developmental Bioengineering Research Grants (EBRG) (R21) CA294038, the Human Frontier Science Program Early-Career Research Grant RGY0088/2021, the Yale Liver Center Pilot Award P30 DK034989, the Yale Lion Heart Pilot Grant, the NIH Director’s Transformative Research Award EB037112, and the DoD CDMRP Ovarian Cancer Research Program Investigator-Initiated Research Award OC240151, HT9425-25-1-0598.

R.V.P was supported by the US National Science Foundation grant MCB-2227268 and the St. Jude Children’s Research Hospital collaborative on the Biology and Biophysics of RNP Granules.

## Contributions

X.S. conceptualized and supervised the project and designed the experiments. Y.X designed, performed the experiments, and analysed data. J.G., K.S., L.Z., Y.T., M.S. and A.A. designed the experiments and contributed to reagent preparation. D.T.T. and A.P. performed the ICP-MS experiment under the supervision of R.V.P. Y.X., X.S. and R.V.P. wrote the manuscript with input from all the other authors.

---

**Extended Data Fig. 1.**
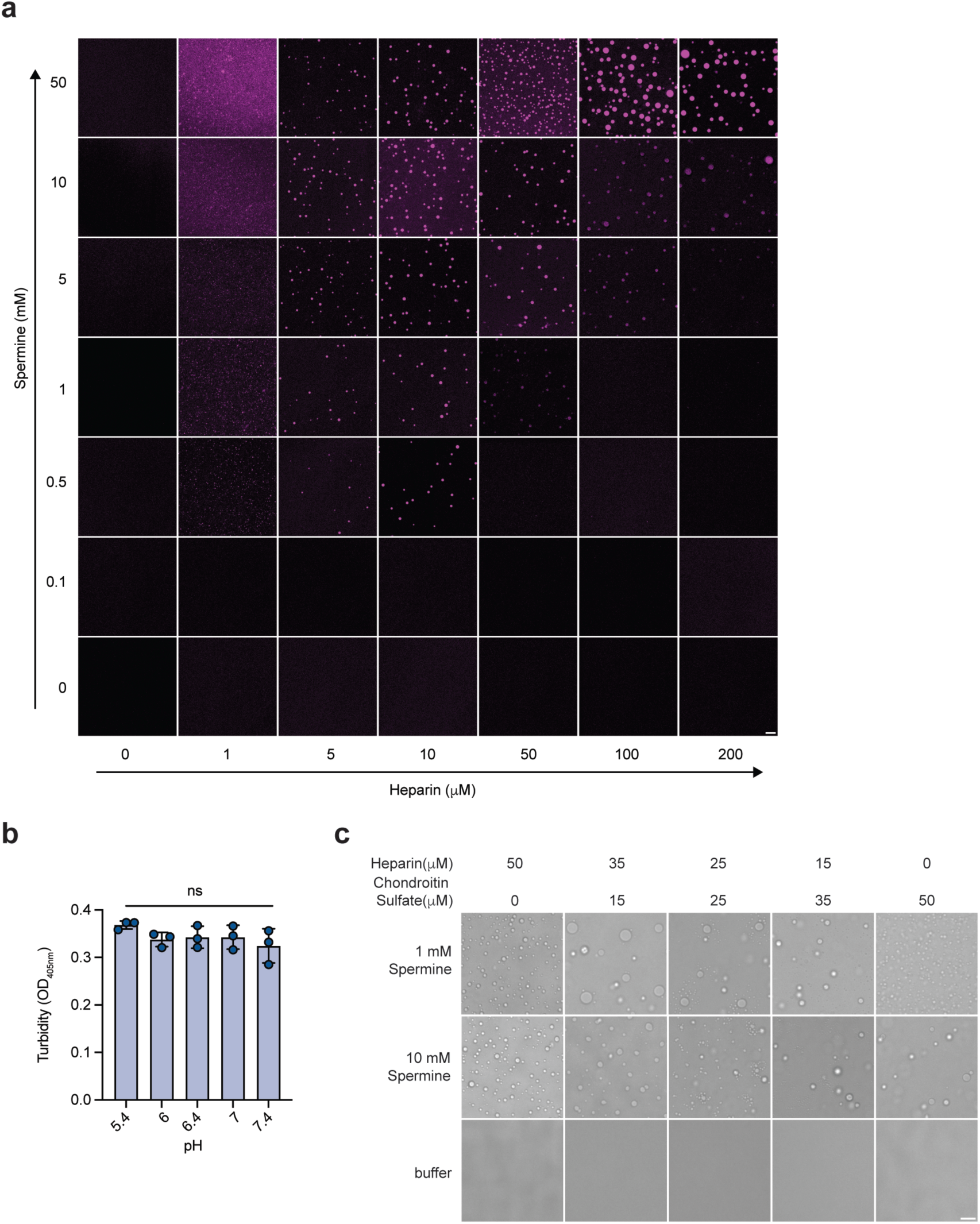
: Polyamines form condensates with heparin. **a.** Fluorescent images of Cy3-labeled heparin in the presence of increasing concentrations of spermine in 50 mM HEPES, pH 7.4 buffer related to Fig. 1c. Scale bar, 10 μm. **b.** Turbidity of heparin-spermine condensate at varying pH. Condensates were assembled by 100 μM heparin and 10 mM spermine in Na_2_HPO_4_-citrate buffer at the indicated pH. N = 3 technical replicates. **c.** Condensation of varying heparin and chondroitin sulfate concentrations in the presence of spermine. Scale bar, 20 μm. Data are presented as Mean ± SD. One representative experiment from 3 independent experiments was shown. ns *p* > 0.05.

**Extended Data Fig. 2.**
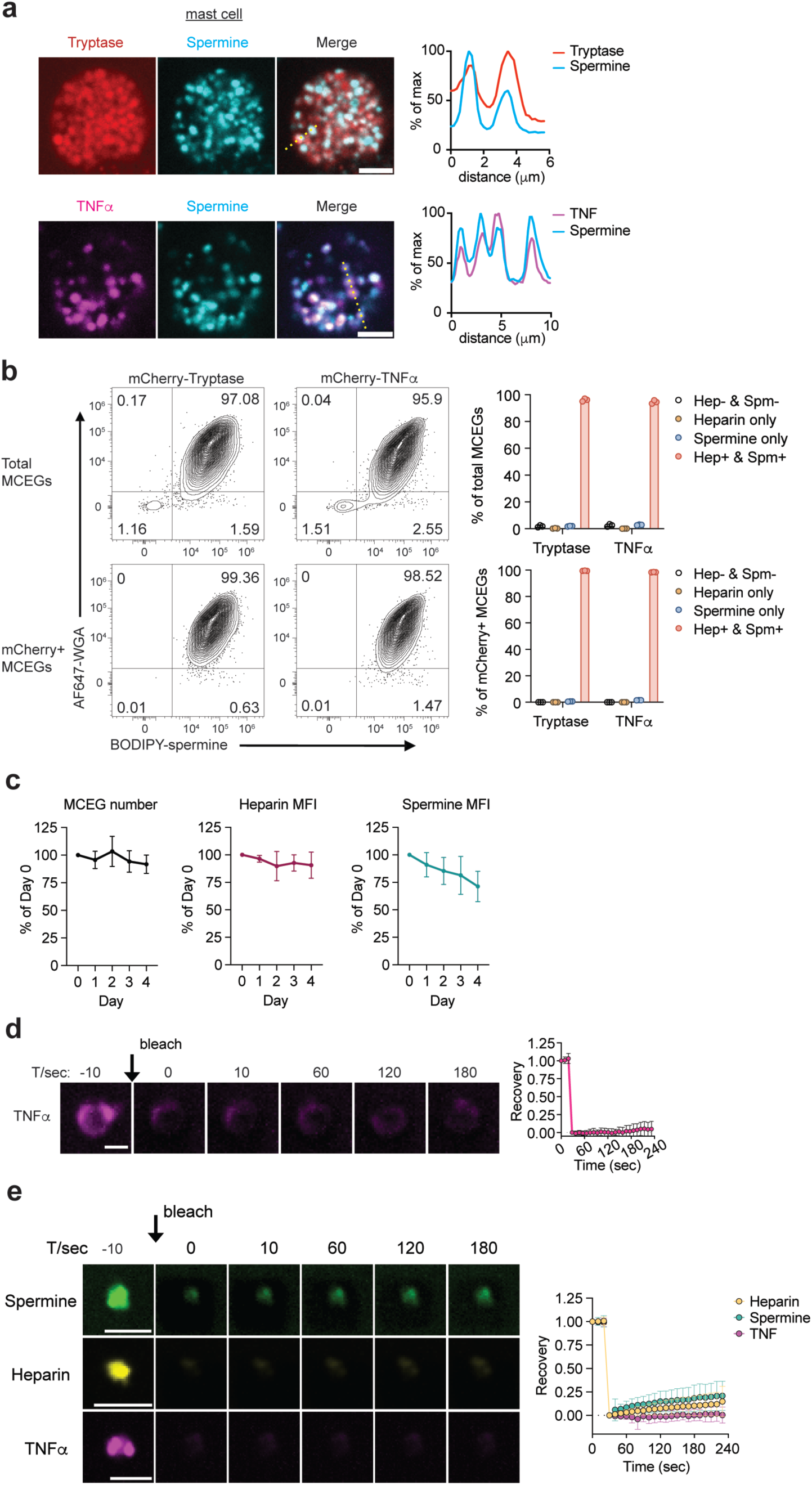
: Spermine is enriched in mast cell extracellular granules (MCEGs). **a.** Fluorescence images of mCherry-Tryptase (red) or mCherry-TNFα (magenta) expressed BMMCs loaded with 20 μM BODIPY-spermine (green) for 1 hr. Right panels, fluorescence intensity of BODIPY-spermine and mCherry-Tryptase or mCherry-TNFα along the yellow dashed line in the merged image. Scale bar, 5 μm. **b.** Analysis of spermine and heparin in MCEGs. Spermine and heparin levels were detected by flow cytometry in MCEGs released from BODIPY-preloaded mast cells expressing mCherry-tryptase or mCherry-TNFα (Left panels). Heparin was labelled with AF647-WGA. The composition analysis showed the proportion of heparin-only, spermine-only, double positive (Hep+ & Spm+), or negative (Hep- & Spm-) MCEGs within the total MCEG population (upper panels) or within mCherry-mediator-retained MCEGs (lower panels). N = 3 replicates. **c.** Stability of MCEGs in heparin- and spermine-free buffers. MCEGs released from BODIPY-spermine-loaded BMMCs were collected after IgE/Ag stimulation by centrifugation. The number of MCEGs and mean fluorescent intensity (MFI) of heparin (labeled with Avidin-sulforhodamine 101) and BODIPY-spermine in MCEGs were determined by flow cytometry following incubation at 37°C in Tyrode’s buffer at the indicated time. Raw counts for each group were normalized to Day 0 values (set as 100%). N = 3 replicates. **d.** FRAP of reconstituted TNFα condensates. Right panel showed the fluorescence recovery of Cy3-labelled TNFα (1.25 μM) in reconstituted condensates formed by 50 μM heparin and 5 mM spermine. N = 23 individual condensates. Scale bar, 2μm. **e.** FRAP of spermine-, heparin-, or TNFα-contained MCEGs. MCEGs released from mast cells pre-loaded with BODIPY-spermine (green), sRd101-avidin (yellow), or expressing mCherry-TNFα (magenta) were separated from cell bodies by low-speed centrifugation, and photobleached in the presence of soluble supernatant following IgE/Ag stimulation. Fluorescence recovery is shown in the right panel. Data represent individual MCEG measurements: N = 28 (sRd101-avidin/heparin), N = 39 (spermine), or N = 14 (TNFα). Scale bar, 5 μm. Data are presented as Mean ± SD. One representative experiment from 3 independent experiments was shown.

**Extended Data Fig. 3.**
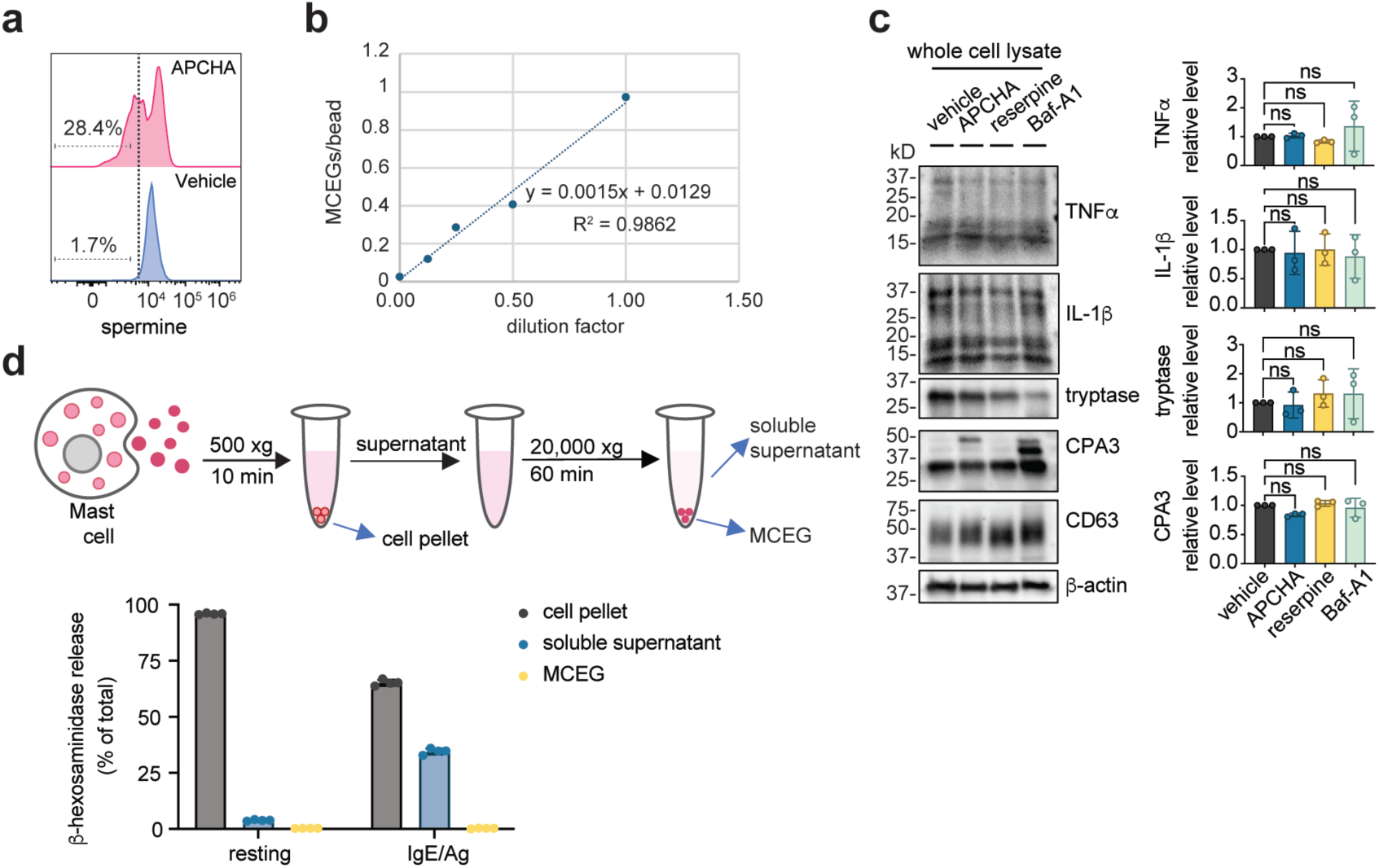
: Depletion of spermine impedes granule assembly. **a.** Spermine levels in APCHA-treated cells were determined by flow cytometry. Dashed line depicts the spermine-positive gate. **b.** Validation of avidin bead-based MCEG quantification method. Serially diluted MCEGs were incubated with equal numbers of avidin beads for 1 hr at room temperature. The number of MCEGs prior to avidin bead incubation was determined by flow cytometry. The number of MCEGs per bead was plotted against the dilution factor in the lower right panel. **c.** Western blot analysis of mast cell mediators in whole cell lysate from inhibitors-treated BMMCs. Band intensities for each mediator were quantified and normalized to β-actin. All normalized values were expressed as fold change relative to the vehicle control and plotted in the right panels (each data point represents one independent experiment, n=3). **d.** Beta-hexosaminidase levels in different cell fractions as separated by centrifugation (upper panel) were expressed as percentages of total β-hexosaminidase in the cells. N = 4 biological replicates. Data are presented as Mean ± SD. One representative of three independent experiments was shown. ns *p* > 0.05.

**Extended Data Fig. 4.**
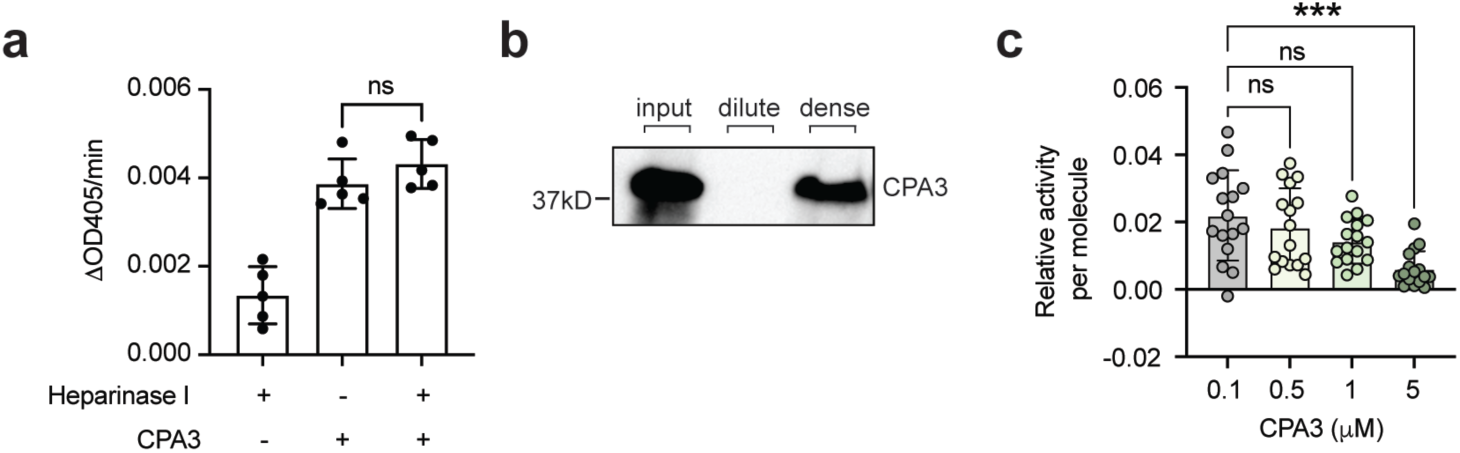
: Heparin-spermine condensates create a metal ion-rich, alkaline environment for enhanced activity of CPA3 **a.** Differences in M-2245 absorbance at 405nm per minute in the presence of heparinase I, CPA3, or a combination of heparinase I and CPA3. N = 5 biological replicates. **b.** Western blot analysis of CPA3 partitioning in the dilute phase and dense phase. **c.** The relative CPA3 protease activity per molecule at the indicated concentrations. N = 16 biological replicates pooled from 4 individual experiments. Data are presented as Mean ± SD. One representative of two (**a**-**b**) or four (**c**) independent experiments was shown. ns *p* > 0.05, *** *p* < 0.001.

**Supplementary Table 1.** : List of identified proteins in the MCEGs of IgE/Ag-stimulated BMMCs.

